# Group or solitary dispersal: worker presence and number favour the success of colony foundation in ants

**DOI:** 10.1101/2023.06.20.545674

**Authors:** Basile Finand, Nicolas Loeuille, Céline Bocquet, Pierre Fédérici, Thibaud Monnin

**Affiliations:** Sorbonne Université, Université Paris Cité, Université Paris Est Créteil, CNRS, INRAE, IRD, Institut d’Ecologie et des Sciences de l’Environnement de Paris (UMR7618), 75005 Paris, France

**Keywords:** Ants, Competition-colonization trade-off, Growth analysis, Survival analysis

## Abstract

Dispersal strategies are highly variable. Any strategy is associated to costs and benefits, and understanding which factors favour or disfavour a strategy is a key issue in ecology and evolution. Ants exhibit different dispersal and colony foundation strategies. Some species have winged queens that disperse solitarily and far by flight, and that found new colonies alone. Others have apterous queens that disperse with workers over short walking distances, and found new colonies as a group (colony fission). The putative benefits conferred by workers have been little studied and quantified, because comparing the costs and benefits of solitary vs. group dispersal and foundation is difficult when comparing different species. We did this using the ant *Myrmecina graminicola*, one of the few species that use both strategies. Young queens were mated and allowed to found new colonies in the laboratory, with either zero, two or four workers. We monitored the survival and growth of foundations over one year. The presence of workers increased both survival and growth, with more workers yielding higher growth. These results show the benefit of dispersing and founding in a group. The presence of few workers (as little as two workers) was sufficient to provide benefits, suggesting group foundation does not require a dramatic decrease in the number of propagules produced in *M. graminicola*. Our results support the hypothesis that the two strategies coexist along a competition-colonization trade-off, where solitary foundation offers a colonization advantage while group foundation has a competitive advantage.

## INTRODUCTION

Dispersal is the movement of individuals that impacts gene flow through space (Ronce 2007). It is a key process in ecology and evolution that affects population dynamics, species distribution, genetic diversity, and local adaptation (Travis et al. 2013). Dispersal is costly yet it is favoured as it allows colonising empty habitats and decreases kin competition (Hamilton and May 1977; Comins et al. 1980). Alternative dispersal strategies differ in costs and benefits, such as when dispersing alone or in a group (also called budding or swarming). Typically, solitary dispersal allows dispersing farther (Cote et al. 2010) but individuals suffer higher predation, body mass loss and stress (Maag et al. 2019). Conversely, group dispersal dilutes predation (Olson et al. 2013) and yields higher establishment success (Memmott et al. 2005; Lange and Marshall 2016), but competition for resources occurs between group members (Lodé et al. 2021). In social insects group dispersal (i.e. queens dispersing with workers) decreases the number of propagules a mother colony can produce compared to solitary dispersal (queens dispersing without workers). Several theoretical and empirical studies show a selection for group dispersal, e.g. when it protects against predation (Holling 1959; Treherne and Foster 1981; Jeschke and Tollrian 2007; Olson et al. 2013; Parthasarathy and Somanathan 2018) or otherwise increases survival (Ridley 2012; Nakamaru et al. 2014), when group members are related (Koykka and Wild 2015), or when group dispersal is sufficient to limit competition with parents (Nakamaru et al. 2014). In addition, it is worth noticing that dispersing solitarily or in a group has a strong impact on the success of settlement in the new habitat, which is the last phase of dispersal.

Ants are good models to study the benefits and costs of solitary versus group dispersal and foundation of new colonies. Firstly, as mentioned above, kin selection can favour group dispersal and ants from a given colony are typically highly related (Hughes et al. 2008; Koykka and Wild 2015). Secondly, there is variability in dispersal and colony founding strategies across species (reviewed in Cronin et al., 2013; Hakala et al., 2019). Some species perform solitary dispersal and foundation (called independent colony foundation) which is the ancestral dispersal strategy (Keller 1991; Heinze and Tsuji 1995; Brown and Bonhoeffer 2003). Their colonies produce many winged queens that disperse solitarily by flying over long distances and found a new colony alone. This strategy is characterised by the production of many propagules (i.e. founding queens), high dispersal, high mortality, and small incipient colonies (the solitary queen). Other species perform group dispersal and foundation (called dependent colony foundation or colony fission). In that case, colonies produce few queens and they are typically unable to fly (apterous or brachypterous) or lack energetic reserves for independent colony foundation (winged microgynes), and they disperse over short distances by walking with a group of workers to found new colonies (Keller and Passera 1989). This strategy has the opposite characteristics of producing fewer propagules (each queen is accompanied by workers), low dispersal, low mortality, and larger incipient colonies. The number of propagules and the dispersal distance of each strategy define its colonising ability and result from queen morphology, whereas survival and size of incipient colonies define its competitive ability and stem from the presence and number of workers. The differences in dispersal and competitiveness between solitary and group foundations, summarised above, suggest that these strategies respond to a colonisation/competition trade-off. Indeed, this trade-off typically produces species that are either good competitor but poor colonizer, or the reverse, and it favours the coexistence of species in meta-communities (Levins and Culver 1971; Tilman 1994; Calcagno et al. 2006). Ant species typically pursue either one strategy or the other, i.e. either solitary or group dispersal and foundation. This makes it difficult to study the costs and benefits of each strategy due to many other confounding factors varying among species. However, some ant species pursue the two strategies (i.e. produce both winged and apterous queens) (Briese 1983; Ohkawara et al. 1993; Heinze 1993; Buschinger and Schreiber 2002) and are thus ideal models for understanding the benefits and costs of each colony foundation strategies, including in terms of survival and growth of incipient colonies.

The presence and number of workers are a main determinant of the survival and growth of colonies, with large colonies surviving better than small ones (Cole et al. 2022). However, very few studies have investigated the effects of workers on newly founded colonies, which are the stage in a colony life cycle that experiences both the highest mortality and the highest growth (Hölldobler and Wilson 1990). Workers increase the survival of *Camponotus yamaokai* queens that found colonies by fission in the laboratory (Shiroto et al. 2011). In the Argentine ant *Linepithema humile*, which naturally founds by colony fission, queens need a hundred workers or more to thrive, and queens founding solitarily in the laboratory do not survive or do not lay eggs (Hee et al. 2000). In *Lasius neglectus*, young queens store less body reserves than in species founding colonies solitarily, yet they can found solitarily in the laboratory, but they produce fewer workers than queens founding with workers (Espadaler and Rey 2001). These studies show the benefit of worker presence, but they used species where the two strategies (solitary foundation vs colony fission) do not occur naturally (ie, without polymorphism). It is expected that providing workers to queens that naturally found solitarily increases their success (e.g. it is used for bumblebees used by the pollinating industry, Gretenkord & Drescher, 1997; Treanore et al., 2021), and that depriving queens that naturally found by colony fission of workers lowers theirs. However, how species that are capable of pursuing both strategies do so remains poorly understood. This likely depends on the costs and benefits conferred by each strategy in the light of the colonisation/competition trade-off (Levins and Culver 1971; Tilman 1994), and here we investigate the survival and growth of colonies founded according to the two strategies.

We use the ant *Myrmecina graminicola* as a model species. It naturally uses colony fission and independent foundation in the same populations (Buschinger et al. 2003). Its queen polymorphism (winged vs apterous) is genetically mediated (Buschinger 2005). We used young queens that emerged and mated in the laboratory to found new colonies with one queen and either no worker or a varied number of workers. We reared these colonies over one year and measured their survival and growth, as a proxy of fitness. We expected the presence of workers to increase colony survival and growth, with more workers yielding larger effects (see references above), and aimed at quantifying this benefit in order to better qualify the costs and benefits each strategy confers in the light of the colonisation/competition trade-off (e.g. allocating more workers to colonies founded by fission trades-off with the number of foundations that can be produced).

## MATERIAL AND METHODS

### Colony sampling and production of sexuals

Colonies were sampled between May and July 2020 and between 11:00 and 16:00 in 16 forests of Ile-de-France near Paris (France) (Figure S1, Table S1). Forests were 17 to 55 km away from Paris, with closest and furthest forest pairs 6.7 to 99.9 km apart, respectively. 102 colonies were found by lifting stones and mosses, under which they are known to occur (Buschinger and Schreiber 2002), and by randomly digging in the ground. They were set up in the laboratory within 48 hours. 22 colonies had no queen, three had apterous queens and 77 had winged (dealated) queens. 76 colonies had one queen (monogyny) and four colonies had several queens (polygyny) (Table S1). Polygynous colonies consisted of one colony with 12 apterous queens and three colonies with several dealated queens each. The reproductive status of queens (mated or virgin, egg-laying or not) was unknown, in particular in polygynous colonies. Hereafter, winged queens refer to queens born with wings, even though they may have cut off their wings and become dealated, and apterous queens refers to queens born without wings.

We reared all queenright colonies until they started producing sexuals in August 2020. Each colony was housed in a 12 x 12 cm box serving as a foraging area and containing a glass-covered nest dug out in a layer of plaster and a water tube (Figure S2). They were fed twice a week (see below). Only colonies with winged queens produced sexuals, and all new queens were of the winged morph. 12 colonies produced queens only, 18 produced males only, and 15 produced both queens and males. In addition, we found five new queens and 54 males that had escaped from their colonies and whose origin could thus not be ascertained. We discarded the new queen escapees but, owing to the scarcity of males, we used the male escapees. Sexual production amounted to a total of 444 queens and 152 males (F:M sex ratio of 2.92) (Table S1). We considered that a male or female was sexually mature when it left the nest for the foraging area and tried to fly.

We used winged queens for both independent colony foundation and colony fission, rather than winged queens for the former and apterous queens for the latter as occurs in nature. This is primarily because we had no young apterous queens to use given that our three source colonies with apterous queens produced no new queens in the laboratory. However, winged and apterous queens of *M. graminicola* are very similar. They have the same size and differ only by the morphology of the thorax, with apterous queens displaying a diversity of thorax morphologies varying from worker-like to almost winged-queen like (Buschinger and Schreiber 2002). Using winged or apterous queens across treatments would have introduced an additional factor that would have needed to be accounted for, e.g. by implementing a crossed experimental design which would have required setting up many additional colony foundations. The use of winged queens only is further explored in the discussion.

### Mating of sexuals

We followed and adapted the protocol of Buschinger (2005) to mate our sexuals. We used mating boxes consisting of a 14.5 x 18.5 cm plastic box coated with fluon, and containing a water tube and two cardboard bridges (1.5 cm width x 7 cm height) to promote flying of sexuals. The sexuals from all our source colonies did not become mature simultaneously. Instead, they matured over several weeks, including within the same colony (Table S2). One mating box was set up for each colony producing new queens. Every day, all new queens and males present in the foraging area of their colonies were collected. Queens were moved to the mating box corresponding to their colony of origin. Males were divided among mating boxes, excluding the one corresponding to their colony of origin. Therefore, each mating box contained queens from one colony and a diversity of males from other colonies. Mating boxes were placed in the sunlight. Some matings were observed but it was impossible to visually confirm that all queens had mated. Queens were thus considered to be mated when they tore their wings off. 21 queens could not be used because they became sexually mature after the last male had died.

### Set up and rearing of colony foundations

We set up 203 colony foundations in the laboratory following three treatments: queen alone to mimic independent colony foundation (n = 72 replicates), or queen assisted by two (n = 65) or four workers (n = 66) to mimic colony fission with different group sizes. Queens and assisting workers were from the same colony. We used two and four workers as relevant group sizes based on data from the closely related Japanese species *M. nipponica*, because propagule size is unknown in *M. graminicola*. *M. nipponica* also performs the two strategies of colony foundation (Ohkawara et al. 1993). It has colonies of similar size than *M. graminicola*, and it allocates on average 2.3 workers (ranging from one to four) to found new colonies by fission (Murakami et al., 2000).

25 of our 203 foundations were removed from the experiment because the queen was inadvertently moved from its box when replacing the water tube or removing the food (i.e. she was lost or moved to another colony). We therefore studied 178 foundations (58, 59 and 61 foundations with zero, two and four workers, respectively). The number of workers assisting the founding queen was held constant. That is, when one worker died, she was replaced by another worker from the same colony of origin.

Foundations were set up by placing the queen, with or without workers, in a 12 x 12 cm rearing box containing a water tube, ad libitum food and a nest (Figure S2). The nest was made up of two 7.6 x 2.5 cm microscope slides separated by 2 mm high walls. It was connected to a 2.5 cm long plaster end that was kept wet by a water tube placed vertically on top of it so as to generate a humidity gradient in the nest. The nest was covered by a piece of black plastic to provide obscurity. Food was provided twice a week during spring, summer and fall and once a week during winter. It consisted of fruit pieces (apple or pear), fruit sauce blended with honey, and dead insects (mealworms or *Drosophila* flies).

Foundations were reared in temperature-controlled chambers (CTS TP 10/600, CTS Gmbh). We reproduced seasons following (Buschinger and Schreiber 2002). Because sexuals emerged over several weeks and matings occurred from the 18^th^ of August 2020 to the 25^th^ of September 2020, foundations were not initiated simultaneously. Therefore, they differed in the amount of time they spent under fall and winter conditions. However, there was no difference across treatment because we were careful to allocate queens to treatments equally over time. We decided to stop winter at the same date for all foundations in order to synchronize them. Foundations therefore spent between 21 and 59 days in the first summer (2020), between 20 and 41 days in fall, between 131 and 152 days in winter, and they all spent 70 days in spring and 122 days in the second summer (2021). The temperatures of fall and spring were 16°C during 12h and 20°C during 12h, 18°C for 10h and 23°C for 14h during summer and 10°C all day during winter.

The experiment lasted about one year. We carried out a weekly census of the number of eggs, larvae, nymphs, and new workers in all foundations under a binocular microscope, and noted whether the queen was alive or dead, and whether she was in the nest or in the foraging area (Figure S2). Queens were dissected at the end of the experiment to confirm that they were mated, including the 49 queens that died during the course of the experiment and which had been preserved in alcohol until dissection. Unfortunately, alcohol altered soft tissues in such a manner that these dissections proved unreliable as the spermatheca was often lost. The 129 queens that survived until the end of the experiment were frozen and rapidly dissected.

### Statistics

Statistical analyses were performed with R 3.6.2 (R Core Team, 2019). We carried out a survival analysis which, necessarily, is based on the death of queens whose dissection was unreliable. Some may have been virgin and incapable of founding a colony, but they would be evenly distributed among the three treatments (as occurs for the virgin queens that survived until the end of the experiment, see results). We used a log-rank test to assess the effect of the treatment (foundation with zero, two or four workers) on queen survival, and compared each pair of treatments with post-hoc Tukey’s test. We used the R packages *survival* and *survminer* (Therneau and Grambsch 2000; Therneau 2022).

We performed a growth analysis considering only the foundations with a mated queen that survived until the end of the experiment. To analyse foundation growth (number of eggs, larvae, nymphs, and new workers) at the end of the experiment depending on the treatment, we used a GLMM with a Poisson error distribution using the R package *lme4* and the R package *emmeans* (Bates et al. 2015). Given that some queens came from the same mother colony, we included the colony of origin of queens as a random effect in the model to account for genetic autocorrelation.

## RESULTS

49 queens died during the course of the experiment and could not be dissected successfully because of alcohol preservation. 129 queens survived until the end of the experiment. Of these, 21 were virgin (Figure S3) and there was no difference in the proportion of virgin queens between the three treatments (5 virgin vs 29 mated, 9 vs 37, and 7 vs 42 for treatments with zero, two and four workers respectively, Chi-square (df = 2, n = 129) = 0.57, p = 0.75).

### Survival analysis

As predicted, the presence of workers increased the survival of the founding queen (Log-rank test Chi-square = 8.1, p = 0.02) (Figure 1). 58.6% (34 out of 58) of solitary queens survived until the end of the experiment against 78% (46 out of 59) of queens with two workers and 80.3% (49 out of 61) of queens with four workers. That is, worker presence increased queen survival by one third. However, when workers were present, there was no significant effect of having more workers as the difference in survival between queens with two and four workers is not statistically significant (Pairwise comparisons: 0 vs 2 workers, p = 0.04 ; 0 vs 4, p = 0.04 ; 2 vs 4, p = 0.8).

**Figure 1:**
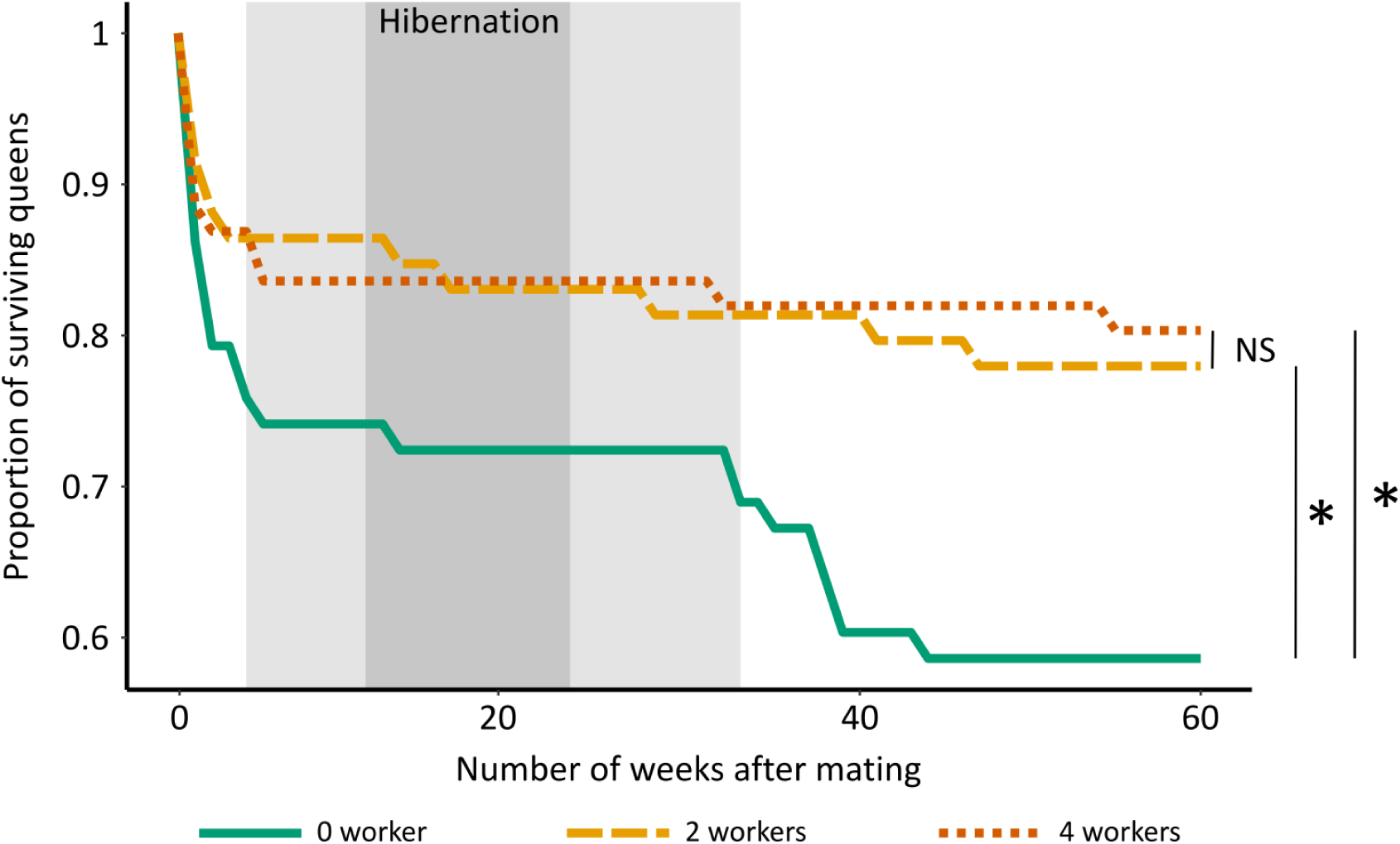
Survival of queens over time. The grey area shows the hibernation period, with light and dark greys denoting periods when only some or all colonies were hibernating, respectively. Queens founding colonies with two or four workers (colony fission, orange lines) survive better than queens founding with no worker (independent foundation, green line, Log-rank test Chi-square = 8.1, p = 0.02). Queens with two workers do not differ in survival from queens with four workers (dark and light orange lines, p = 0.8).

The temporal dynamic shows four phases of mortality (Figure 1). First, an initial period of high queen mortality at the beginning of the experiment when foundations were set up. This mortality differs among treatments. The proportion of queens alive drops to 86% when workers are present and to 74% for solitary queens. Second, a period of low mortality during hibernation. Third, a second period of high queen mortality right after hibernation. It affects solitary queens only, so that the proportion of solitary queens alive drops to 59%. Fourth, another period of low queen mortality until the cessation of the experiment.

The weekly censuses of queen presence in the nest revealed that queens with two and four workers rapidly settled in the nest following the onset of the experiment, whereas solitary queens wandered for longer in the foraging area (Figure 2). Surprisingly, most solitary queens resumed wandering in the foraging area after the hibernation phase, while few of the queens with workers did (Figure 2). After a few weeks, these solitary queens eventually also settled in the nest.

**Figure 2:**
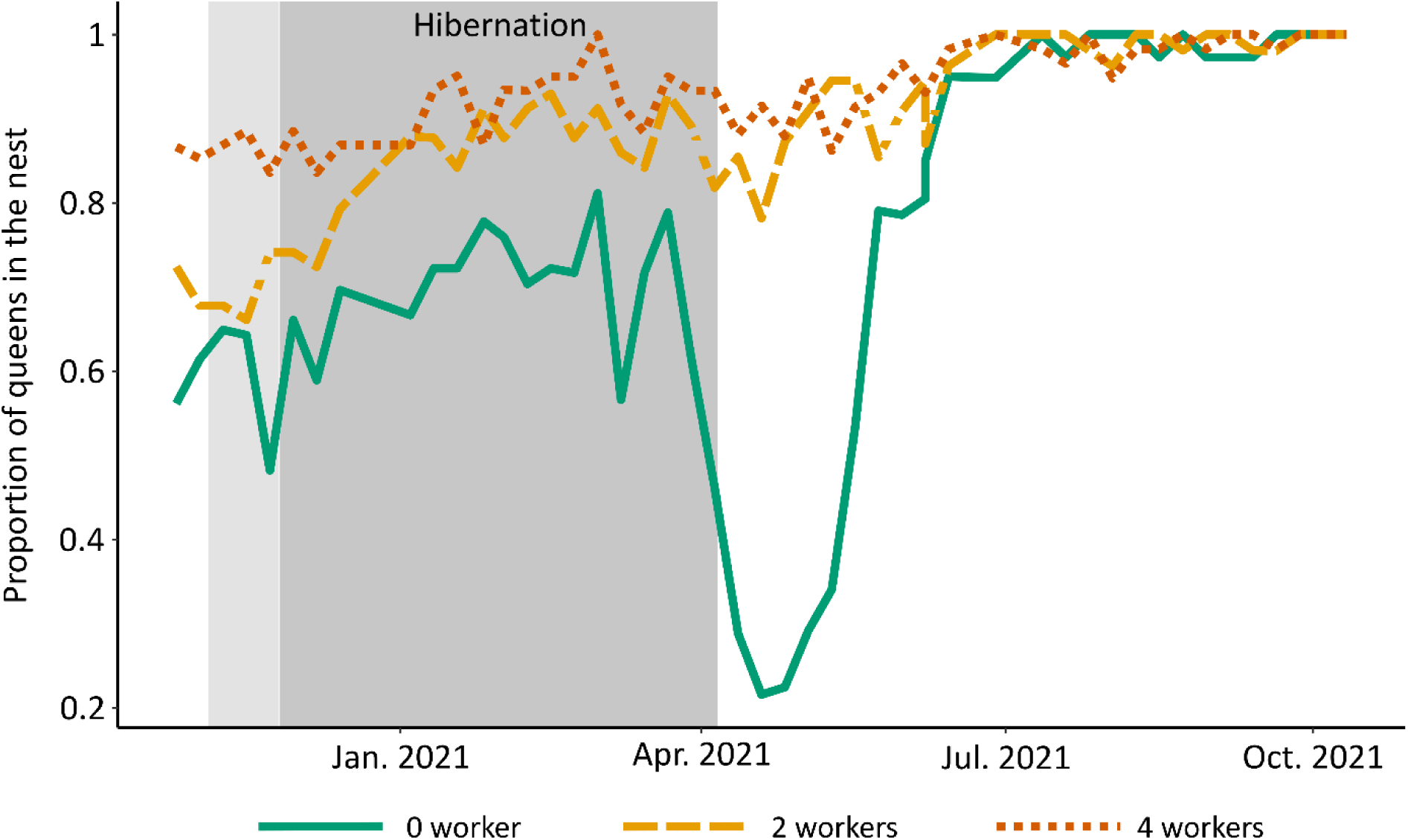
Presence of queens in the nest over time. The x-axis is synchronised across colonies by the date of end of hibernation. The grey area shows the hibernation period, with light and dark greys denoting periods when only some or all colonies were hibernating, respectively. Solitary founding queens leave the nest for the foraging area after hibernation, unlike queens with workers. Note that the presence of queens in their nest was not initially recorded, that is the start of the x-axis does not correspond to the start of the experiment.

### Growth analysis

As intuitively expected, the presence of workers favoured the growth of the incipient colonies (Figure 3a). There are several major differences across treatments. First, queens founding their colony with two or four workers started laying eggs almost immediately after colony foundation and before hibernation (Figure 3c,d), while solitary queens started laying eggs after hibernation only (Figure 3b). When workers were present, their number affected queen fertility as the number of eggs and larvae was higher in the treatment with four workers than in the treatment with two workers (3.07 +/- 3.04 items vs 1.57 +/- 2.38 items at the pre-hibernation peak in brood number, on the 12^th^ of October 2020).

**Figure 3:**
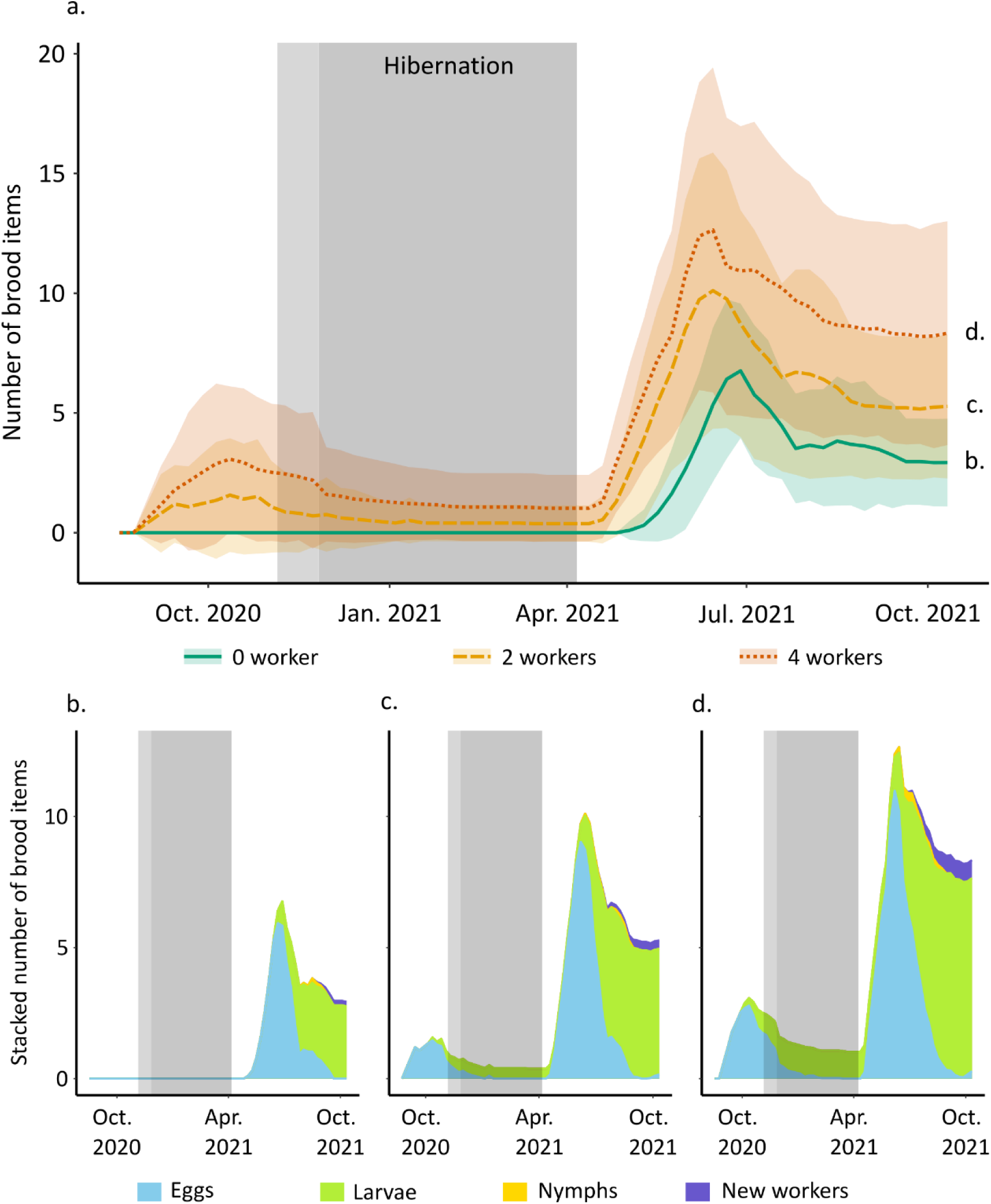
Colony growth over time. The x-axis is synchronised across colonies by the date of end of hibernation. The grey area shows the hibernation period, with light and dark greys denoting periods when only some or all colonies were hibernating, respectively. (a) temporal dynamic of the number of brood items (sum of eggs, larvae, nymphs and new workers) for each treatment (mean+/- SD). Queens with workers produce brood before hibernation whereas solitary queens do not, and colony growth increases with the number of workers. (b-d) stacked

Second, after hibernation, queens founding with workers resumed egg laying two weeks earlier than solitary queens (on the 19^th^ of April 2021 vs 3^rd^ of May 2021, Figure 3b vs c,d). The presence and number of workers affected colony growth. The numbers of brood items produced differ among the three treatments and increase with worker number. The production of queens with zero, two and four workers peaked at 6.76 +/- 2.81, 10.11 +/- 5.77 and 12.64 +/- 6.79 brood items, respectively (Figure 3a). Because of the delayed onset of egg-laying by queens without workers, the number of brood items they produced peaked two weeks later (28^th^ of June 2021) than for queens with two or four workers (14^th^ of June 2021 for both).

Third, there is a difference between treatments in the number of brood items present at the end of the experiment (GLMM, Chi-square = 88.284, p < 0.0001) (Figure 4). The presence of workers increases the productivity of colony foundations, with more workers yielding higher productivity. Queens with zero, two and four workers had 2.93 +/- 1.83, 5.27 +/- 3.01 and 8.33 +/- 4.67 brood items at the end of the experiment, respectively (pairwise tests: 0 vs 2 workers: p < 0.0001 ; 0 vs 4: p < 0.0001 ; 2 vs 4 workers: p < 0.0001). That is, worker presence increased brood production by 80% (two workers) or 185% (four workers). Most of these items were larvae (Figure 3). Few larvae developed in nymphs, and all nymphs became workers.

**Figure 4:**
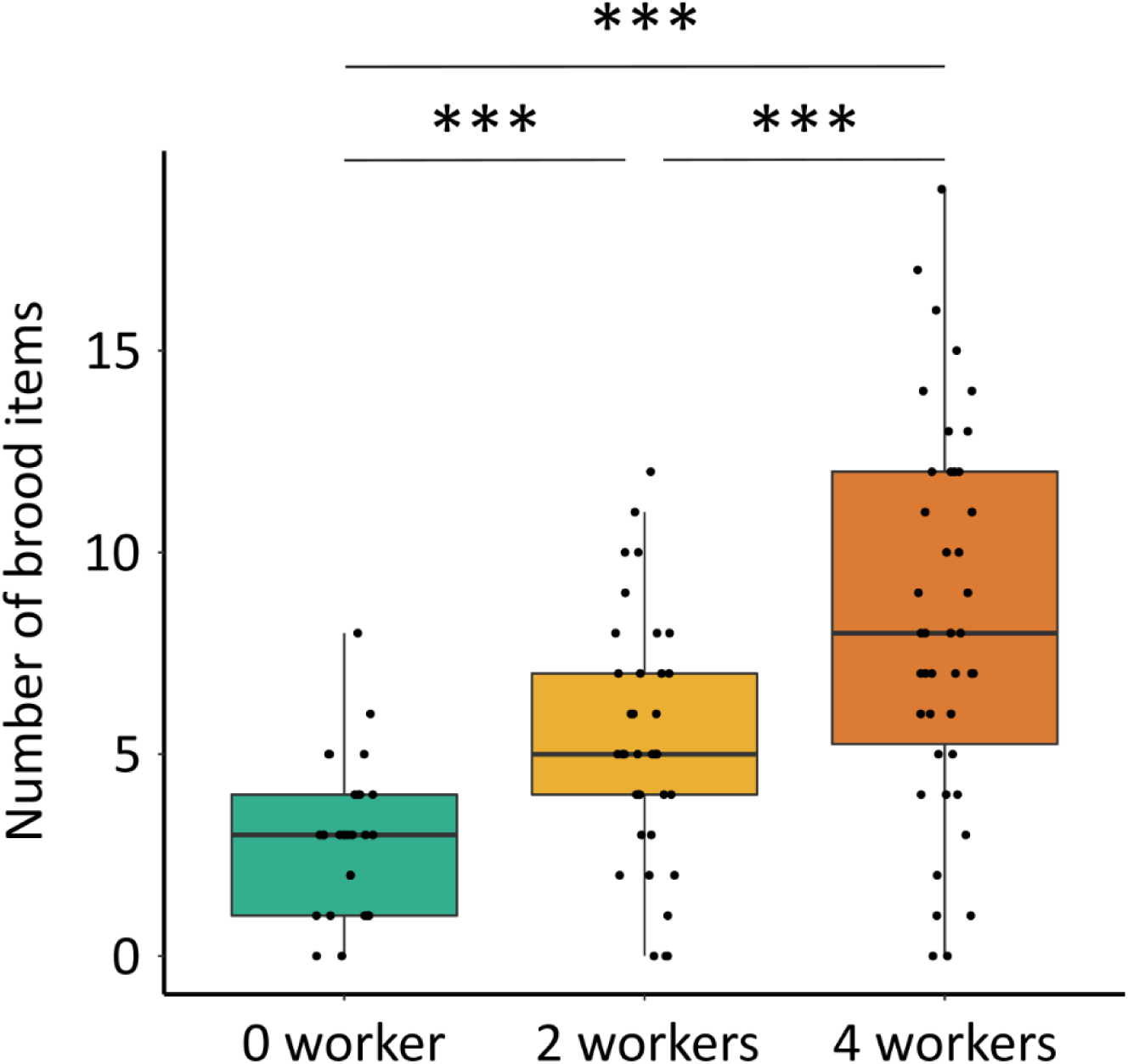
Colony growth (sum of eggs, larvae, nymphs and new workers) at the end of the experiment increases with the number of workers (GLMM, Chi-square = 88.284, p < 0.0001).

## DISCUSSION

This study is, to our knowledge, the first to demonstrate and quantify the benefits conferred by founding colonies with workers compared to solitarily in an ant species that uses these two strategies. Using a species that is polymorphic in terms of its colony foundation strategy gives unique insights on the conditions that favour either strategy and the maintenance of the polymorphism. It shows that colonies founded with worker had higher survival (increased by one third) and growth (increased by 80 to 185%) than those founded solitarily, and that founding with a larger group of workers provided larger benefits.

Our experimental design was constrained because our source colonies did not produce apterous queens. Therefore, we used winged queens to set up both solitary- and group-founded colonies, whereas the latter are normally founded by apterous queens in nature. This is unlikely to have affected our findings. In *M. graminicola*, winged and apterous queens differ in wing morphology and eyes size but are otherwise similar in body size including abdomen size (Buschinger and Schreiber 2002). They have similar fertility hence using winged queens in lieu of apterous queens should not affect the results, provided they interact appropriately with workers. This was clearly the case as they adjusted their behaviour to the presence of workers, by modifying their exploratory behaviour and timing of egg-laying. Whether this behavioural plasticity of winged queens could occur in nature is unclear.

Our findings of a higher survival of group foundations in the laboratory is likely conservative. Queen and incipient colony mortalities are much higher in nature than in our laboratory experiment. For example, only 16.3% of *Atta texana* queens survive after ninety days (Marti et al. 2015), 6% of *Solenopsis invicta* colonies after fifty days (Tschinkel 1992) and 7.6% of *Crematogaster ashmeadi* colonies after one year (Hahn and Tschinkel 1997). These high mortalities result from predation and environmental hazards during dispersal and, once the first workers start foraging, competition with larger pre-established colonies. Solitary founding queens suffer more from these factors than group-founding queens (e.g. Cronin et al., 2013) so that the differential in survival between our colony foundation modalities is likely higher under natural conditions.

The beneficial effect of worker presence was expected. They carry out all tasks required by the colony except egg laying (e.g. brood rearing, nest building and maintenance, foraging for food, nest cleaning and protection against pathogens, nest defence, Hölldobler & Wilson, 1990) and bear the associated costs, thereby alleviating the burden of the queen. However, few studies have assessed the benefits of the presence and of the number of workers on the growth of newly founded colonies (see introduction). Our findings agree with the facts that species that found colonies by fission produce brood faster than species that found colonies by solitarily queens (Keller and Passera 1990), and that, in the fissioning Argentine ant *Linepithema humile*, larger groups of workers yield larger increases in brood production, until a threshold number of workers above which growth reaches a plateau (Hee et al. 2000).

The Japanese congeneric species *M. nipponica* also produces both winged and apterous queens (Cronin et al., 2020). Apterous queens are more abundant in cold environments, suggesting that the advantages of colony foundation by colony fission are particularly important in these challenging environments. This may be because cold temperatures are physiologically stressful so that solitary queens may suffer higher mortality than queens hibernating in incipient colonies founded by fission (Cronin et al. 2020). In addition, the short warm season may restrict colony growth, and this would be more detrimental to smaller colonies (Cronin et al. 2016). Our results on *M. graminicola* are in line with this. Queens founding with workers had lower mortality and higher growth than solitary-founding queens under laboratory conditions, and in the Paris region apterous queens are more abundant in city parks than in forests (Finand et al. 2023), which may be because city parks are a challenging environment (e. g. heavy metal pollution) and/or because high habitat fragmentation selects against dispersal by flight in cities.

Theoretical works have suggested that species may coexist along a trade-off between colonization and competitive abilities (Levins and Culver 1971; Tilman 1994). Species then differ in colonization and in competitive abilities, with a trade-off between the two, so that coloniser species can move to sites that competitive species cannot reach. The importance of this trade-off among species has been highlighted in various groups (eg, ciliates, Cadotte et al., 2006, or passerine birds, Rodríguez et al., 2007). Our results support the idea that this trade-off not only occurs between species, but also occurs within species, here between the two queen morphs of *M. graminicola*. They show that colonies founded by a queen with workers have higher survival and growth than colonies founded by a solitary queen. That is, there is a competitive benefit of dispersing in a group (as happens for apterous queens in nature), while solitary queens rely on a colonizing strategy (high dispersal distance, high mortality). The competition-colonization trade-off can foster the maintenance of diversity among species (Tilman 1994), and we propose that when it occurs within species it helps maintaining polymorphism. Polymorphism in reproductive strategies allows coping with contrasted environments and is especially expected in spatially structured environments (Parvinen 2002; Parvinen et al. 2020; Finand et al. 2022). It has been observed in various empirical systems, for instance in the weed *Crepis sancta* where dispersal is restricted in fragmented cities (Cheptou et al. 2008) and in various ant species where group foundation predominates in cold environments (*M. nipponica*, Cronin et al., 2020), in patchy environments (*L. canadensis,* Heinze, 1993) or in periods of drought (*Chelaner sp.,* Briese, 1983).

The higher growth rate of colonies founded with workers allows them to reach maturity sooner, so that this strategy locally outcompetes solitary foundation and dominates the environment (Cronin et al. 2016). In *M. graminicola*, as little as two workers are sufficient to increase colony survival and growth in the laboratory, in agreement with the observed size of incipient colonies in *M. nipponica* (Murakami et al. 2000). This suggests that colony fission in these species entails a relatively low restriction in the number of propagules that can be produced compared to solitary foundations. Indeed, queens being only slightly larger than workers, and other things being equal, a colony producing apterous queens that found colonies with two workers each (i.e. propagules of three individuals) may presumably produce one third as many propagules as a colony producing winged solitary-founding queens. The source colonies that we had collected to produce sexuals in the laboratory had 53 +/- 17 (mean +/- SD) workers (Table S1). This suggests that propagules of three individuals represent 6% of colony size. This relatively low cost of fissioning propagules may allow these species to produce more propagules than many of the species that reproduce by colony fission. Indeed, and although data are lacking for most species, many fissioning species appear to produce only one (Gotwald 1995) or very few propagules (Chéron et al. 2011; Doums et al. 2013; Villalta et al. 2015).

Our finding that foundations with workers survive and grow better than those without workers parallels the higher survival and growth of colonies founded by several queens (pleometrosis) compared to those founded by a solitary queen (haplometrosis) in some ant species displaying a polymorphism in this respect (Tschinkel and Howard 1983; Rissing and Pollock 1991; Ostwald et al. 2021). In both cases, the larger initial group size is the determinant of the higher growth rate, and larger foundations outgrow and outcompete smaller ones through scramble competition for resources or through aggressive behaviours such as territoriality or brood raiding depending on species’ biology. However, fissioning propagules are a permanent association between one queen and related workers whereas pleometrosis is often a transitory association between unrelated queens with, typically, all but one queen being killed by other queens and/or workers after the emergence of the first workers (Sommer & Hölldobler, 1995). In addition, fissioning propagules have restricted dispersal, unlike pleometrotic queens. Pleometrosis also occurs in some facultatively social bees, where it provides benefits by group-enhanced defence against predators and prevention of brood failure and can result in permanent polygyny (Schwarz et al., 1998).

Our results are congruent with theoretical works that predict the emergence of group dispersal when it confers higher survival (Nakamaru et al. 2014), dilutes the risk of predation (Olson et al. 2013) or when group members are highly related (Koykka and Wild 2015). Indeed, group dispersers experience lower mortality or higher fitness in various taxonomic groups such as spiders (Parthasarathy and Somanathan 2018), Arabian gazelles (Ridley 2012), tadpole toad (Watt et al. 1997) and *Daphnia* (Pijanowska and Kowalczewski 1997). Larger group size is also beneficial in several social mammals where it increases offspring production and survival, such as mongooses (Rood, 1990), black-backed jackals (Moehlman 1979), red foxes (Macdonald 1979) or primates (Majolo et al. 2008). In some bird species, juveniles stay with their parents for a time to help raise their siblings. This increases their survival and development but also permits a larger clutch size (Reyer 1984; Arnold and Owens 1998; Ridley 2007).

During our experiment, we observed a phase of relatively high mortality in the first days after mating, at the onset of colony foundation. This likely reflects the energetic cost and stress of mating and finding a suitable nesting site. Queens that founded with workers immediately settled in the nest and started laying eggs. In contrast, queens that founded solitarily delayed egg laying to after the hibernation, which slowed their colony growth compared to that of queens founding with workers. In addition, many solitary-founding queens explored the foraging arena following hibernation, before eventually settling in the nest and producing eggs. This correlated with a second wave of mortality that was specific to solitary-founding queens, presumably because this exploration incurs energetic costs and stresses (e.g. exposure to desiccation) but a causal link is yet to be established. Why solitary queens were active in the foraging area is unclear. One possibility is that they were seeking a suitable nesting site, another that they had consumed all their body reserves during hibernation and foraged for food. Few larvae developed into workers during our one-year long experiment, hence colony growth consisted almost solely in the production of eggs and larvae (Figure 3). This agrees with the fact that *M. graminicola* queens produce few larvae during the first year of foundation (Buschinger 2005). Similar low growth during the first year of colony foundation have been documented in other ant species such as *Acromyrmex octopinosus* (three to seven workers produced in several months, Fernandez-Marin et al., 2003) and *Mycocepurus smithii* (one worker in nine months, Fernández-Marín et al., 2005).

Our results are consistent with the hypothesis that a colonization/competition trade-off occurs intra-specifically and this may help maintain queen polymorphism in *M. graminicola*. Further theoretical and empirical studies are necessary to understand the evolution of colony foundation strategies. Field studies may provide valuable information on how environmental factors (habitat fragmentation, aggregation, and quality) affect the two dispersal and foundation strategies in terms of queen morphology and propagule size and number, which would provide a better understanding of their evolution and occurrence within the colonization-competition trade-off framework.

## Supporting information

SI_article

## DECLARATIONS

### Ethics approval

Not applicable

### Consent to participate

Not applicable

### Consent for publication

Not applicable

### Availability of data and material

If the paper is accepted, the data will be available on zenodo.

### Code availability

If the paper is accepted, the code for analysis will be available on GitHub.

### Authors’ contributions

BF, NL and TM conceived the ideas and designed methodology; BF collected the data, with help from CB and PF; BF analyzed the data; BF, NL and TM wrote the manuscript. All authors gave final approval for publication.

