## Supplementary material for "Group or solitary dispersal: worker presence and number favour the success of colony foundation in ants": SI_article

**SUPPLEMENTARY INFORMATIONS**

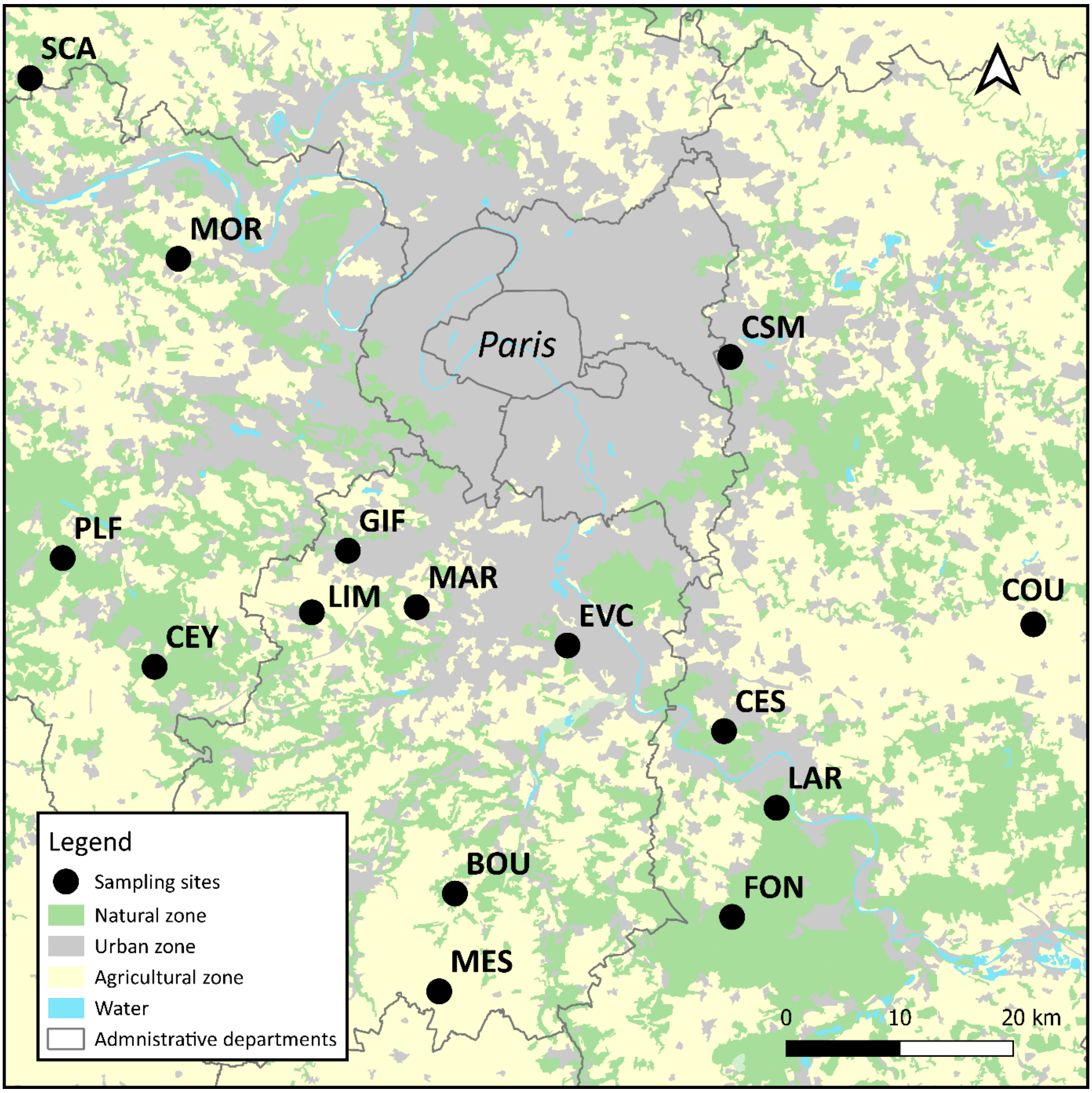


**Figure S1**: Map of the sampling sites

BOU: Bouville (48.4221 2.26205), CES: Cesson (48.55474 2.5899), CEY: Clairefontaine-en-Yvelines (48.60292 1.90214), COU: Courpalay (48.64094 2.95876), CSM: Champs-sur-Marne (48.85005 2.59058), EVC: Evry-Courcouronnes (48.62142 2.40127), FON: Fontainebleau (48.41563 2.62169), GAM: Gambais (48.75785 1.71486), GIF: Gif-sur-Yvettes (48.69484

2.13738), LAR: La Rochette (48.49384, 2.65326), LIM: Limours (48.64728 2.08862), MAR: Marcoussis (48.65913 2.21452), MES: Mespuits (48.36013 2.23541), MOR: Morainvilliers (48.92478 1.93579), PLF: Poigny-la-forêt (48.694206 1.799004), SCA: Saint-Cyr-en-Arthies (49.06572 1.74926)

(Map source: UE-SDES, CORINE Land Cover, 2012)

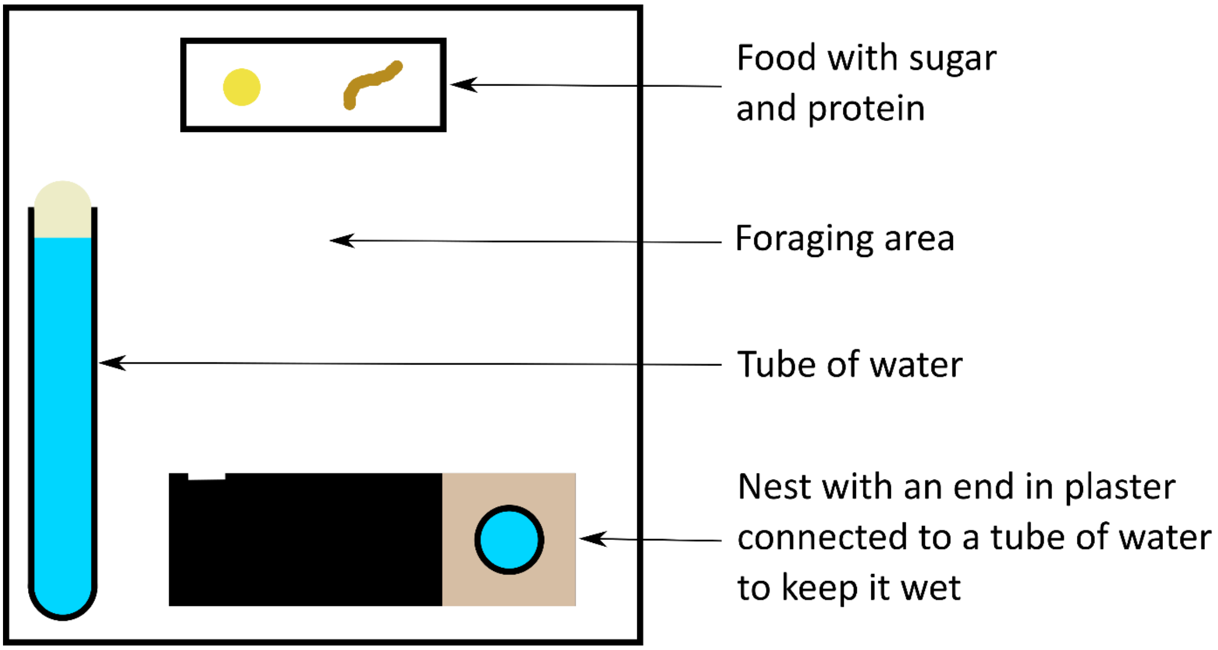


**Figure S2:** Representation of the rearing box

a) b)

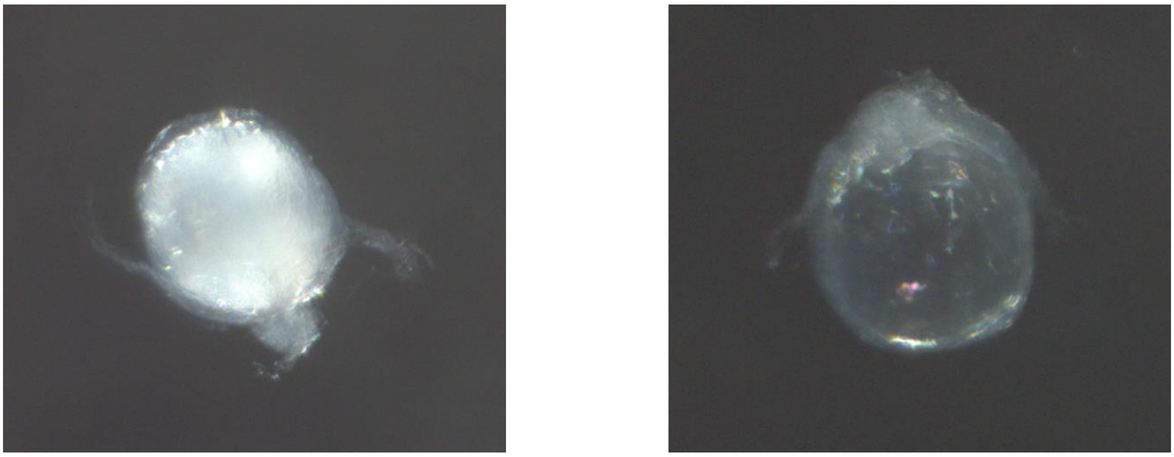

**Figure S3:** Examples of spermatheca of (a) a mated queen and (b) a virgin queen

**Table S1:** Characteristics of colonies used for the production of sexuals. The last line corresponds to sexuals that have escaped form the mother colony and were wandering in the chambers. Therefore, we cannot link them to a mother colony.

BOU: Bouville (48.4221 2.26205), CES: Cesson (48.55474 2.5899), CEY: Clairefontaine-en-Yvelines (48.60292 1.90214), COU: Courpalay (48.64094 2.95876), CSM: Champs-sur-Marne (48.85005 2.59058), EVC: Evry-Courcouronnes (48.62142 2.40127), FON: Fontainebleau (48.41563 2.62169), GAM: Gambais (48.75785 1.71486), GIF: Gif-sur-Yvettes (48.69484 2.13738), LAR: La Rochette (48.49384, 2.65326), LIM: Limours (48.64728 2.08862), MAR: Marcoussis (48.65913 2.21452), MES: Mespuits (48.36013 2.23541), MOR: Morainvilliers (48.92478 1.93579), PLF: Poigny-la-forêt (48.694206 1.799004), SCA: Saint-Cyr-en-Arthies (49.06572 1.74926)

| **Colony** | **Locality** | **Number of winged dealated queens** | **Number of apterous queens** | **Number of new queens produced** | **Number of males produced** | **Number of new queens mated** |
|---|---|---|---|---|---|---|
| 38 | LAR | 1 | 0 | 11 | 0 | 1 |
| 39 | LAR | 0 | 0 | 0 | 0 | 0 |
| 40 | LAR | 1 | 0 | 0 | 0 | 0 |
| 41 | LAR | 1 | 0 | 0 | 0 | 0 |
| 42 | LAR | 1 | 0 | 2 | 3 | 0 |
| 43 | LAR | 1 | 0 | 0 | 0 | 0 |
| 44 | LAR | 1 | 0 | 13 | 5 | 9 |

| | | | | | | |
|---|---|---|---|---|---|---|
| 45 | LAR | 1 | 0 | 10 | 0 | 5 |
| 46 | LAR | 1 | 0 | 0 | 0 | 0 |
| 47 | LAR | 2 | 0 | 9 | 0 | 8 |
| 48 | LAR | 1 | 0 | 0 | 0 | 0 |
| 49 | LAR | 0 | 0 | 0 | 0 | 0 |
| 50 | LAR | 0 | 12 | 0 | 0 | 0 |
| 51 | LAR | 0 | 0 | 0 | 0 | 0 |
| 52 | LAR | 1 | 0 | 1 | 0 | 0 |
| 53 | LAR | 0 | 0 | 12 | 12 | 7 |
| 54 | LAR | 1 | 0 | 0 | 0 | 0 |
| 55 | LAR | 0 | 0 | 0 | 0 | 0 |
| 56 | LAR | 1 | 0 | 6 | 1 | 0 |
| 57 | LAR | 0 | 1 | 0 | 0 | 0 |
| 58 | LAR | 1 | 0 | 0 | 0 | 0 |
| 59 | COU | 1 | 0 | 34 | 1 | 19 |
| 60 | EVC | 1 | 0 | 0 | 0 | 0 |
| 61 | EVC | 1 | 0 | 12 | 0 | 7 |
| 62 | EVC | 1 | 0 | 0 | 0 | 0 |
| 63 | EVC | 0 | 0 | 0 | 0 | 0 |
| 64 | CEY | 1 | 0 | 0 | 0 | 0 |
| 65 | CEY | 1 | 0 | 0 | 0 | 0 |
| 66 | CEY | 1 | 0 | 31 | 4 | 20 |
| 67 | CEY | 1 | 0 | 31 | 1 | 15 |
| 68 | CEY | 0 | 0 | 0 | 0 | 0 |

| | | | | | | |
|---|---|---|---|---|---|---|
| 69 | CEY | 0 | 0 | 1 | 1 | 0 |
| 70 | CEY | 1 | 0 | 0 | 0 | 0 |
| 71 | CEY | 1 | 0 | 15 | 2 | 1 |
| 72 | CEY | 1 | 0 | 48 | 0 | 23 |
| 73 | CEY | 1 | 0 | 19 | 1 | 6 |
| 74 | CEY | 1 | 0 | 21 | 4 | 12 |
| 75 | CEY | 1 | 0 | 30 | 1 | 17 |
| 76 | CEY | 1 | 0 | 0 | 0 | 0 |
| 77 | CEY | 1 | 0 | 0 | 0 | 0 |
| 78 | CEY | 1 | 0 | 4 | 0 | 0 |
| 79 | CEY | 0 | 0 | 2 | 0 | 0 |
| 80 | CEY | 1 | 0 | 7 | 0 | 4 |
| 81 | CEY | 1 | 0 | 0 | 0 | 0 |
| 82 | CEY | 1 | 0 | 0 | 1 | 0 |
| 83 | SCA | 1 | 0 | 0 | 5 | 0 |
| 84 | MES | 1 | 0 | 0 | 0 | 0 |
| 85 | MES | 1 | 0 | 10 | 16 | 7 |
| 86 | GIF | 1 | 0 | 0 | 0 | 0 |
| 87 | GIF | 1 | 0 | 0 | 0 | 0 |
| 88 | GIF | 0 | 0 | 0 | 0 | 0 |
| 89 | GIF | 1 | 0 | 0 | 3 | 0 |
| 90 | GIF | 1 | 0 | 0 | 2 | 0 |
| 91 | GIF | 0 | 0 | 0 | 1 | 0 |
| 92 | GIF | 1 | 0 | 0 | 0 | 0 |

| | | | | | | |
|---|---|---|---|---|---|---|
| 93 | GIF | 1 | 0 | 10 | 3 | 4 |
| 94 | GIF | 0 | 0 | 0 | 2 | 0 |
| 95 | GIF | 1 | 0 | 0 | 0 | 0 |
| 96 | GIF | 1 | 0 | 0 | 1 | 0 |
| 97 | GIF | 0 | 0 | 0 | 4 | 0 |
| 98 | LIM | 1 | 0 | 4 | 1 | 3 |
| 99 | LIM | 1 | 0 | 0 | 0 | 0 |
| 100 | LIM | 1 | 0 | 0 | 0 | 0 |
| 101 | CSM | 1 | 0 | 0 | 0 | 0 |
| 102 | CSM | 1 | 0 | 17 | 0 | 10 |
| 103 | CSM | 1 | 0 | 0 | 0 | 0 |
| 104 | CES | 0 | 0 | 0 | 0 | 0 |
| 105 | CES | 1 | 0 | 8 | 0 | 1 |
| 106 | CES | 1 | 0 | 0 | 0 | 0 |
| 107 | CES | 0 | 0 | 27 | 3 | 9 |
| 108 | MOR | 1 | 0 | 0 | 0 | 0 |
| 109 | MOR | 1 | 0 | 3 | 0 | 2 |
| 110 | MOR | 0 | 0 | 0 | 0 | 0 |
| 111 | MOR | 1 | 0 | 5 | 2 | 5 |
| 112 | MOR | 1 | 0 | 0 | 0 | 0 |
| 113 | MOR | 1 | 0 | 0 | 3 | 0 |
| 114 | MOR | 1 | 0 | 0 | 0 | 0 |
| 115 | MOR | 1 | 0 | 4 | 0 | 2 |
| 116 | MOR | 1 | 0 | 0 | 0 | 0 |

| | | | | | | |
|---|---|---|---|---|---|---|
| 117 | MOR | 0 | 1 | 0 | 0 | 0 |
| 118 | MOR | 1 | 0 | 0 | 0 | 0 |
| 119 | MOR | 1 | 0 | 0 | 2 | 1 |
| 120 | MOR | 0 | 0 | 0 | 0 | 0 |
| 121 | MOR | 1 | 0 | 0 | 0 | 0 |
| 122 | MOR | 1 | 0 | 0 | 0 | 0 |
| 123 | MOR | 1 | 0 | 0 | 0 | 0 |
| 124 | MAR | 1 | 0 | 0 | 0 | 0 |
| 125 | MAR | 1 | 0 | 2 | 8 | 0 |
| 126 | MAR | 1 | 0 | 3 | 0 | 2 |
| 127 | MAR | 0 | 0 | 0 | 0 | 0 |
| 128 | MAR | 1 | 0 | 0 | 0 | 0 |
| 129 | MAR | 2 | 0 | 0 | 1 | 0 |
| 130 | MAR | 1 | 0 | 12 | 0 | 0 |
| 131 | MAR | 1 | 0 | 0 | 0 | 0 |
| 132 | GAM | 7 | 0 | 2 | 1 | 0 |
| 133 | GAM | 1 | 0 | 2 | 1 | 0 |
| 134 | FON | 0 | 0 | 0 | 0 | 0 |
| 135 | PLF | 0 | 0 | 0 | 0 | 0 |
| 136 | BOU | 0 | 0 | 3 | 1 | 0 |
| 137 | BOU | 1 | 0 | 0 | 1 | 0 |
| 138 | BOU | 0 | 0 | 7 | 0 | 3 |
| 139 | BOU | 1 | 0 | 1 | 0 | 0 |
| Unknown | Unknown | NA | NA | 5 | 54 | NA |

**Table S2:** Dates of transfer of sexuals in mating boxes.

| Colony | Sex | Number | Date |
|---|---|---|---|
| 45 | Queen | 3 | 04 August 2020 |
| 53 | Male | 11 | 04 August 2020 |
| 44 | Male | 3 | 04 August 2020 |
| 38 | Queen | 3 | 04 August 2020 |
| 56 | Queen | 4 | 04 August 2020 |
| 42 | Male | 3 | 04 August 2020 |
| 38 | Queen | 7 | 18 August 2020 |
| 74 | Queen | 7 | 18 August 2020 |
| 45 | Queen | 7 | 18 August 2020 |
| 47 | Queen | 5 | 18 August 2020 |
| 44 | Queen | 5 | 18 August 2020 |
| Wandering male | Male | 20 | 18 August 2020 |
| 44 | Male | 1 | 19 August 2020 |
| 59 | Queen | 22 | 19 August 2020 |
| 85 | Male | 12 | 19 August 2020 |
| 72 | Queen | 9 | 19 August 2020 |
| 75 | Queen | 12 | 19 August 2020 |
| 83 | Male | 4 | 19 August 2020 |
| 44 | Queen | 3 | 19 August 2020 |
| 47 | Queen | 4 | 19 August 2020 |
| 66 | Male | 2 | 19 August 2020 |

| | | | |
|---|---|---|---|
| 67 | Male | 1 | 19 August 2020 |
| 74 | Male | 2 | 19 August 2020 |
| 61 | Queen | 6 | 19 August 2020 |
| 113 | Male | 1 | 19 August 2020 |
| 67 | Queen | 7 | 19 August 2020 |
| Wandering male | Male | 1 | 20 August 2020 |
| 38 | Queen | 1 | 20 August 2020 |
| 44 | Queen | 4 | 20 August 2020 |
| 85 | Queen | 7 | 20 August 2020 |
| 85 | Male | 1 | 20 August 2020 |
| 113 | Male | 2 | 20 August 2020 |
| 71 | Queen | 2 | 20 August 2020 |
| 72 | Queen | 22 | 20 August 2020 |
| 73 | Queen | 6 | 20 August 2020 |
| 73 | Male | 1 | 20 August 2020 |
| 74 | Queen | 14 | 20 August 2020 |
| 74 | Male | 1 | 20 August 2020 |
| 75 | Queen | 12 | 20 August 2020 |
| 78 | Queen | 2 | 20 August 2020 |
| 53 | Queen | 5 | 20 August 2020 |
| 56 | Queen | 1 | 20 August 2020 |
| 59 | Queen | 9 | 20 August 2020 |
| 61 | Queen | 6 | 20 August 2020 |
| 67 | Queen | 4 | 20 August 2020 |

| | | | |
|---|---|---|---|
| 69 | Male | 1 | 20 August 2020 |
| 66 | Queen | 6 | 20 August 2020 |
| 66 | Male | 1 | 20 August 2020 |
| 97 | Male | 2 | 20 August 2020 |
| Wandering male | Male | 4 | 20 August 2020 |
| 42 | Queen | 2 | 21 August 2020 |
| 44 | Queen | 1 | 21 August 2020 |
| 44 | Male | 1 | 21 August 2020 |
| 52 | Queen | 1 | 21 August 2020 |
| 53 | Queen | 6 | 21 August 2020 |
| 59 | Queen | 3 | 21 August 2020 |
| 59 | Male | 1 | 21 August 2020 |
| 66 | Queen | 22 | 21 August 2020 |
| 67 | Queen | 11 | 21 August 2020 |
| 71 | Queen | 9 | 21 August 2020 |
| 72 | Queen | 14 | 21 August 2020 |
| 73 | Queen | 4 | 21 August 2020 |
| 74 | Male | 1 | 21 August 2020 |
| 75 | Queen | 5 | 21 August 2020 |
| 75 | Male | 1 | 21 August 2020 |
| 78 | Queen | 2 | 21 August 2020 |
| 80 | Queen | 6 | 21 August 2020 |
| 85 | Queen | 2 | 21 August 2020 |
| 85 | Male | 3 | 21 August 2020 |

| | | | |
|---|---|---|---|
| 90 | Male | 1 | 21 August 2020 |
| 102 | Queen | 3 | 21 August 2020 |
| 111 | Queen | 1 | 21 August 2020 |
| 111 | Male | 1 | 21 August 2020 |
| 56 | Queen | 1 | 21 August 2020 |
| 69 | Queen | 1 | 21 August 2020 |
| 125 | Male | 1 | 21 August 2020 |
| 107 | Male | 1 | 21 August 2020 |
| 56 | Male | 1 | 24 August 2020 |
| 67 | Queen | 1 | 24 August 2020 |
| 67 | Queen | 1 | 24 August 2020 |
| 71 | Queen | 1 | 24 August 2020 |
| 72 | Queen | 2 | 24 August 2020 |
| 73 | Queen | 6 | 24 August 2020 |
| 89 | Male | 1 | 24 August 2020 |
| 93 | Queen | 3 | 24 August 2020 |
| 66 | Male | 1 | 24 August 2020 |
| 105 | Queen | 2 | 24 August 2020 |
| 107 | Queen | 1 | 24 August 2020 |
| 125 | Male | 1 | 24 August 2020 |
| 132 | Male | 1 | 24 August 2020 |
| 53 | Queen | 1 | 25 August 2020 |
| 66 | Queen | 2 | 25 August 2020 |
| 71 | Queen | 2 | 25 August 2020 |

| | | | |
|---|---|---|---|
| 71 | Male | 1 | 25 August 2020 |
| 89 | Male | 2 | 25 August 2020 |
| 83 | Male | 1 | 25 August 2020 |
| 85 | Queen | 1 | 25 August 2020 |
| 90 | Male | 1 | 25 August 2020 |
| 93 | Queen | 4 | 25 August 2020 |
| 93 | Male | 2 | 25 August 2020 |
| 97 | Male | 2 | 25 August 2020 |
| 98 | Queen | 4 | 25 August 2020 |
| 98 | Male | 1 | 25 August 2020 |
| 82 | Male | 1 | 25 August 2020 |
| 80 | Queen | 1 | 25 August 2020 |
| 119 | Male | 2 | 25 August 2020 |
| 102 | Queen | 11 | 25 August 2020 |
| 107 | Queen | 11 | 25 August 2020 |
| 107 | Male | 1 | 25 August 2020 |
| 105 | Queen | 3 | 26 August 2020 |
| 130 | Queen | 2 | 26 August 2020 |
| 79 | Queen | 1 | 26 August 2020 |
| 107 | Queen | 3 | 26 August 2020 |
| 129 | Male | 1 | 26 August 2020 |
| Wandering male | Male | 1 | 26 August 2020 |
| Wandering male | Male | 1 | 27 August 2020 |
| 102 | Queen | 2 | 27 August 2020 |

| | | | |
|---|---|---|---|
| 126 | Queen | 3 | 27 August 2020 |
| 130 | Queen | 4 | 27 August 2020 |
| 75 | Queen | 1 | 27 August 2020 |
| 71 | Male | 1 | 27 August 2020 |
| 96 | Male | 1 | 27 August 2020 |
| 111 | Queen | 2 | 27 August 2020 |
| 111 | Male | 1 | 27 August 2020 |
| 107 | Queen | 4 | 27 August 2020 |
| 94 | Male | 1 | 27 August 2020 |
| 125 | Male | 3 | 27 August 2020 |
| 105 | Queen | 1 | 27 August 2020 |
| 115 | Queen | 4 | 27 August 2020 |
| 73 | Queen | 1 | 27 August 2020 |
| 71 | Queen | 1 | 28 August 2020 |
| 107 | Queen | 4 | 28 August 2020 |
| 107 | Male | 1 | 28 August 2020 |
| 109 | Queen | 3 | 28 August 2020 |
| Wandering male | Male | 1 | 28 September 2020 |
| 67 | Queen | 2 | 28 August 2020 |
| 93 | Queen | 3 | 28 August 2020 |
| 93 | Male | 1 | 28 August 2020 |
| 79 | Queen | 1 | 28 August 2020 |
| 72 | Queen | 1 | 28 August 2020 |
| 73 | Queen | 2 | 28 August 2020 |

| | | | |
|---|---|---|---|
| 125 | Queen | 2 | 28 August 2020 |
| 125 | Male | 3 | 28 August 2020 |
| 66 | Queen | 1 | 28 August 2020 |
| 53 | Male | 1 | 02 September 2020 |
| 91 | Male | 1 | 02 September 2020 |
| 105 | Queen | 2 | 02 September 2020 |
| 111 | Queen | 1 | 02 September 2020 |
| 130 | Queen | 1 | 02 September 2020 |
| 133 | Male | 1 | 02 September 2020 |
| 136 | Queen | 1 | 02 September 2020 |
| 137 | Male | 1 | 02 September 2020 |
| 67 | Queen | 4 | 08 September 2020 |
| 130 | Queen | 2 | 08 September 2020 |
| 136 | Queen | 1 | 08 September 2020 |
| 107 | Queen | 1 | 08 September 2020 |
| 138 | Queen | 3 | 08 September 2020 |
| 111 | Queen | 1 | 09 September 2020 |
| 107 | Queen | 2 | 09 September 2020 |
| 132 | Queen | 1 | 09 September 2020 |
| 138 | Queen | 2 | 09 September 2020 |
| 133 | Queen | 1 | 09 September 2020 |
| 102 | Queen | 1 | 09 September 2020 |
| 107 | Queen | 1 | 10 September 2020 |
| 130 | Queen | 2 | 10 September 2020 |

| | | | |
|---|---|---|---|
| 136 | Queen | 1 | 10 September 2020 |
| 67 | Queen | 1 | 10 September 2020 |
| 139 | Queen | 1 | 10 September 2020 |
| 94 | Male | 1 | 11 September 2020 |
| 132 | Queen | 1 | 11 September 2020 |
| 138 | Queen | 1 | 11 September 2020 |
| 138 | Queen | 1 | 15 September 2020 |
| 136 | Male | 1 | 15 September 2020 |
| 133 | Queen | 1 | 15 September 2020 |
| 130 | Queen | 1 | 16 September 2020 |
| Wandering male (Found dead at the end of the sexual production) | Male | 26 | 16 September 2020 |
| Wandering queen (Found dead at the end of the sexual production) | Queen | 5 | 16 September 2020 |
